# Gene regulation inference from single-cell RNA-seq data with linear differential equations and velocity inference

**DOI:** 10.1101/464479

**Authors:** Pierre-Cyril Aubin-Frankowski, Jean-Philippe Vert

## Abstract

Single-cell RNA sequencing (scRNA-seq) offers new possibilities to infer gene regulation networks (GRN) for biological processes involving a notion of time, such as cell differentiation or cell cycles. It also raises many challenges due to the destructive measurements inherent to the technology. In this work we propose a new method named GRISLI for *de novo* GRN inference from scRNA-seq data. GRISLI infers a velocity vector field in the space of scRNA-seq data from profiles of individual data, and models the dynamics of cell trajectories with a linear ordinary differential equation to reconstruct the underlying GRN with a sparse regression procedure. We show on real data that GRISLI outperforms a recently proposed state-of-the-art method for GRN reconstruction from scRNA-seq data.

## 1 Introduction

Single-cell RNA sequencing (scRNA-seq) enables to observe genome-wide cellular activities at the single cell resolution (Kolodziejczyk *et al*., 2015), generating extraordinary expectations for biologists and bringing forth new computational and mathematical challenges. By allowing us to study cell-to-cell variability, scRNA-seq has quickly become a technique of choice to systematically identify cell types in complex samples (Trapnell, 2015; Zeisel *et al*., 2015; Tasic *et al*., 2016) and understand dynamic biological processes such as embryo development (Deng *et al*., 2014), cell differentiation (Lönnberg *et al*., 2017) and cancer (Patel *et al*., 2014).

A fascinating perspective offered by scRNA-seq studies is to understand how genes interact and regulate each other. In particular, by observing how gene expression varies among similar cells subject to stochastic fluctuations or involved in a dynamical process such as differentiation or cell cycle, one may be able to capture statistical or dynamical dependencies between genes which may in turn allow to reverse-engineer a gene regulatory network (GRN) to describe biologically which transcription factors (TF) regulate which genes. While numerous algorithms have been proposed to infer GRN from bulk transcriptomic profiles (e.g., Marbach *et al*., 2012, and references therein), scRNA-seq data raises new opportunities and challenges. On the one hand, the quantity of cells in scRNA-seq studies is often several-fold larger than the number of samples in bulk transcriptomic studies, offering increased statistical power to capture regulatory interactions, and allowing to capture subtle changes in dynamical process. On the other hand, scRNA-seq data are subject to various sources of variability (Kharchenko *et al*., 2014; Risso *et al*., 2018), and the precise type or state of each cell in a population must usually be inferred themselves from the data. In particular, in the case of dynamical processes such as differentiation or cell cycles, several methods have been proposed to automatically infer a *pseudo-time* associated to each individual cell, as reviewed by Cannoodt *et al*. (2016).

As in bulk transcriptomics studies, putative functional interactions between genes can be detected by simple correlation analysis (Moignard *et al*., 2013; Stegle *et al*., 2015; Bacher and Kendziorski, 2016), or through more advanced strategies to capture statistical dependency between genes tailored to scRNA-seq data (Chan *et al*., 2017; Filippi and Holmes, 2017). Aibar *et al*. (2017) refined the detection of gene modules by combining sequence information. However, such statistical associations do not necessarily capture regulatory relationships, which typically require perturbations or temporal experiments to be detected. As scRNA-seq studies can provide an ordering of cells involved in a dynamical process, through experimental time and/or inferred pseudo-time, they offer a unique opportunity to infer regulatory relationships by taking into account the (pseudo-)time information to compare gene expression profiles. For example, Herbach *et al*. (2017) propose a realistic, albeit complex, stochastic dynamical system to model scRNA-seq data, which is only tested on simulated data for networks of two genes due to its computational complexity. Moignard *et al*. (2015) present a formalism to infer a boolean network from single cell qRT-PCR data, but requires to discretize gene expression values to an on/off status in each cell. Ocone *et al*. (2015) propose to infer a GRN by estimating an ordinary differential equation (ODE) from pseudo-time-ordered scRNA-seq data; however, due to the computational complexity of the model selection procedure, the final GRN is limited to be a refinement of a coarse GRN inferred with GENIE3 (Huynh-Thu *et al*., 2010), a method for bulk gene expression. A similar, linear ODE-based formalism was proposed by Matsumoto *et al*. (2017), who designed a more efficient procedure named SCODE to directly infer *de novo* the GRN from scRNA-seq data. SCODE assumes that all cells are on the same trajectory, and estimates the parameters of the ODE by integrating it and optimizing the fit between the integrated model and each individual cell’s transcriptome. However, the resulting optimization problem is computationally intractable, and is solved only approximately by restricting the class of GRN models.

In this work, we follow the same linear ODE-based formalism as SCODE for GRN inference from scRNA-seq data, and propose a new approach, which we name GRISLI, to estimate the parameters of the model. GRISLI first estimates the *velocity* of each cell, i.e., how each gene’s expression is increasing or decreasing in the dynamical process for each cell, and then estimates the structure of the GRN by solving a sparse regression problem to relate the gene expression of a cell to its velocity profile. We solve the sparse regression problem with a variant of stability selection (Meinshausen and Bühlmann, 2010) proposed in TIGRESS (Haury *et al*., 2012), a method for GRN inference from bulk transcriptomics where no velocity is involved since samples are assumed to be near steady state. In spite of a similar ODE formalism, GRISLI differs from SCODE in several aspects: (i) while SCODE assumes that all cells are on the same trajectory, in GRISLI we consider *bundles* of trajectories derived from a large number of initial conditions and do not integrate the ODE; (ii) while SCODE integrates the ODE, leading to a computationally intractable optimization problem to infer the parameters, we solve a simple, convex regression problem that allows us to make no restrictive assumption on the GRN structure and leads to a fast algorithm. These benefits come at the cost of estimating the velocity of each cell, for which we propose a novel procedure based on weighted averages of finite differences with other cells at nearby positions in space-time.

We empirically assess the performance of GRISLI on human and murine scRNA-seq data and show that it outperforms TIGRESS, highlighting the benefits of the ODE-based framework for scRNA-seq data, as well as outperforming SCODE, confirming the relevance of our new estimation procedure.

## 2 Methods

### 2.1 Setting and notations

We consider the problem of inferring a GRN from a set of *C* single-cell transcriptomic profiles *x*_1_,…, *x_C_* ∈ ℝ^*G*^, where *x_i_* ∈ ℝ^*G*^ represents the expression, for the *i*-th cell, of *G* genes. We furthermore assume that the cells are involved in a dynamical process, such as differentiation or cell cycle, and that for each cell *i* ∈ [1, *C*] we have an estimate of a time label *t_i_* ∈ ℝ that describes where the cell is in the process. The time-label *t_i_* is assigned to the *i*-th cell based either on the real experimental time, or on a calculated pseudo-time. Hence we assume given a collection of time-labeled vectors { (*x_i_, t_i_*) ∈ ℝ^*G*^ × ℝ: *i* = 1,…, *C*.}

We model the dynamical process of the cell expression *x*(*t*) ∈ R^*G*^ as a linear ordinary differential equation (ODE) of the form

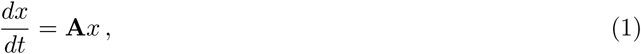

where **A** ∈ ℝ^*G×G*^ characterizes how each gene’s expression level influences the expression dynamics of other genes. Assuming that each gene is regulated by only a few TFs, we assume that **A** is sparse, in the sense that **A***_ij_* ≠ 0 means that the expression of gene *j* influences that of gene *i*, i.e., that gene *j* regulates gene *i*.

Inferring the GRN thus amounts to estimating which entries in **A** are non-zero. To do so, we propose a two-step approach called GRISLI: first we estimate the velocity of each cell *v_i_* = *dx_i_/dt* with an estimator 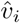, and second we infer non-zero elements of **A** by estimating the support of the regression model (1) from the sample 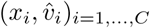 with a stability selection procedure. We detail each step in turn below.

### 2.2 Velocity inference

Given the set of time-labeled vectors (*x_i_, t_i_*) ∈ ℝ^*G*^ × ℝ: *i* = 1,…, *C*, we estimate the velocity 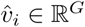 of each cell *i* ∈ [1, *C*] as follows. We first observe that from any other cell (*x_j_, t_j_*), with *t_j_* ≠ *t_i_*, we may form the following velocity estimate based on finite difference:

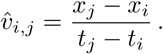

This estimate is interesting only when (i) *t_j_* is not too far from *t_i_*, so that a finite difference is a good approximation of the derivative, and (ii) the trajectories of cells *j* and *i* are close to each other, in the sense that if we were able to observe cell *j* at time *t_i_* then it should be close to *x_i_*. Based on these considerations, we form the velocity estimate 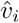 as a weighted average of the 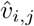, with weights defined by a spatio-temporal kernel *K*(*x, t, x*′, *t*) that quantifies how we believe (*x*′, *t*′) is useful to estimate the velocity at (*x, t*). However, as the points living in the past or the future of a given point (*x_i_, t_i_*) act differently on the velocity, we separate their contributions into two weighted averages.

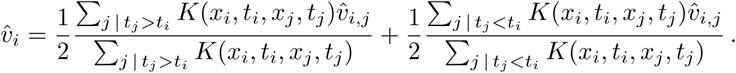

As for the spatio-temporal kernel *K*, we arbitrarily take the following:

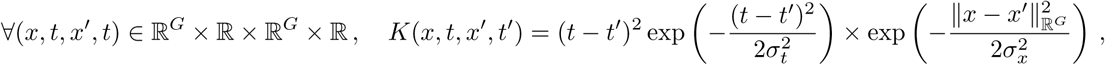

where *σ_x_* and *σ_t_* are fixed respectively to the square root of the 10th percentile of the distribution of distances in space, *x* (resp. in time, *t*).

### 2.3 GRN inference

Once we form the estimate 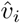 for the velocity *v_i_* = *dx_i_/dt* of each cell *i* = 1,…, *C*, we estimate the GRN by considering (1) as a sparse regression problem of the form 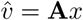, with observations 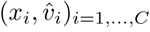 We estimate the non-zero entries of **A** using a stability selection procedure for sparse regression (Meinshausen and Bühlmann, 2010; Haury *et al*., 2012). More precisely, for each candidate regulator *j* ∈ [1, *C*] and target gene *i* ∈ [1, *C*], we compute a score *s*(*i, j*) ∈ (0, 1) which increases when we believe that **A***_ij_* ≠ 0, i.e., that *j* regulates *i*. The score *s*(*i, j*) itself depends on three parameters *R, L* ∈ ℕ and *α* ∈ [0, 1], and is computed through the procedure described by Haury *et al*. (2012), which we now summarize. Let us denote by **X**:= (*x*_1_,…, *x_C_*) ∈ ℝ^*G×C*^ the expression matrix and 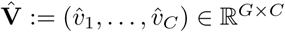 the matrix of estimated velocities. We repeat *R* times a procedure where we create a new expression matrix 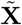 and a new velocity matrix 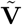 obtained from **X** and 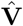 by (i) randomly subsampling ⌊*C/*2⌋ columns (i.e., cells) simultaneously from **X** and 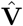, and (ii) multiplying each row *i* of **X** by a different random number *β_i_* uniformly sampled between *α* and 1. We then estimate for every 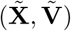 a sparse matrix **A** by solving a lasso regression problem:

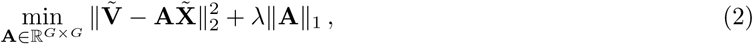

over a grid of regularization parameters *λ* ensuring that we have solutions having from 0 to at least *L* nonzero entries in each row of **A**. For each pair of genes (*i, j*) and each integer *l* ∈ [1, *L*] we measure, among the *R* repeats, the frequency *F* (*i, j, l*) at which **A***_ij_* is nonzero for the solution of (2) when *λ* is set such that *l* entries are nonzero in the *i*-th row of **A**, i.e., when the *j*-th TF is among the top *l* TFs in the regularization path to explain the expression of the *i*-th gene. We then consider the *area score* proposed by Haury *et al*. (2012):

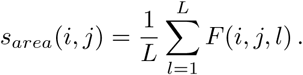

Alternatively, we can consider the original stability selection score proposed by Meinshausen and Bühlmann (2010):

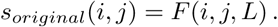

Haury *et al*. (2012) discusses the differences between both scores, and suggests to prefer the area score which is therefore our preferred choice.

The choice of the three parameters *R, L* and *α* is a difficult question. While *R* should typically be as large as possible to reduce random fluctuations of the algorithm, *α* should typically be chosen in the range [0.2, 0.8] according to Haury *et al*. (2012) and *L* should be tested on a large grid of values. In the experiments below we provide results for different values of these parameters to demonstrate the potential of the method. In other applications where some interactions are known, we suggest as well to test predictions over a grid of values, and to pick the model that best matches known interactions.

### 2.4 Data

In order to test GRISLI, we use the two datasets provided online by Matsumoto *et al*. (2017). These are scRNA-Seq datasets where the TF expression is considered as the log-transform of the transcripts per million reads (TPM) or the log-transform of the fragments per millions of kilobases mapped (FPKM). The TF data comes from the RIKEN mouse TFdb for mouse (Kanamori *et al*., 2004) and animalTFDB for human (Hu *et al*., 2018), which we downloaded from the Transcription Factor Regulatory Network database (http://www.regulatorynetworks.org). In each case we kept only the 100 TFs with the highest variance in our datasets to infer the network, as they are the most likely to have an influence over differentiation.

The first dataset, published by Treutlein *et al*. (2016), (named *Data2* in SCODE) comes from a direct reprogramming of murine embryonic fibroblast cells to myocytes at days 0, 2, 5 and 22. This dataset contained 373 cells.

The second dataset (named *Data3* in SCODE) comes from Chu *et al*. (2016) and measures the differentiation of human ES cells to definitive endoderm cells, taken at 0, 12, 24, 36, 72 and 96 h. This dataset contained 758 cells.

### 2.5 Performance evaluation

We evaluate the performance of GRISLI by its area under the receiver operating characteristic curve (AUC), calculated by comparing the predictions of GRISLI (a score for each pair of TFs) to the gold standard regulatory networks. For the sake of comparison, we follow the choices made by Matsumoto *et al*. (2017): we compute the AUC ignoring self-loops (the diagonal elements) and the TFs that do not have an edge in the true network. After discarding the TFs with a variance among the top 100 for which no edge exists in the regulation network taken from the literature, only a smaller number of TFs remains for which we can compute the ROC. There are 40 TFs left for the murine dataset and 49 TFs for the human dataset.

## 3 Results

### 3.1 GRISLI

We propose a new method for Gene Regulatory network Inference from scRNA-seq data with LInear differential equations (GRISLI). The input to GRISLI is a set of time-stamped scRNA-seq data (*x_i_, t_i_*)_*i*=1,…,*C*_, where *C* is the number of cells, *x_i_* is the vector of gene expression for the *i*-th cell and *t_i_* is the time associated to the *i*-th cell; this time can be based either on the real experimental time, or on a calculated pseudo-time. GRISLI combines the dynamical model of SCODE (Matsumoto *et al*., 2017) with the statistical procedure for network estimation of TIGRESS (Haury *et al*., 2012). More precisely, like SCODE we model the dynamics of gene expression as a linear differential equation *dx/dt* = **A***x*, where **A** is sparse and encodes the GRN in its non-zero entries. By integrating this equation, and assuming all cells have the same (unknown) stage *x*_0_, Matsumoto *et al*. (2017) propose to estimate **A** by solving:

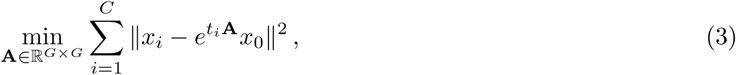

which is however a computationally intractable non-convex optimization problem in **A**; to overcome the difficulty, Matsumoto *et al*. (2017) restrict themselves to low-rank diagonalizable matrices of the form **A** = **WBW**^+^, where **B** is diagonal of small rank, and use further assumptions and heuristics to obtain a tractable algorithm SCODE to optimize successively **W** and **B**.

GRISLI is based on the same dynamical model as SCODE, but exploits it differently. Instead of integrating the dynamical model *dx/dt* = **A***x* as in (3), we see it as a regression problem of the form *v* = **A***x* where *v* = *dx/dt* is the velocity of each cell and take a two-step approach to estimate **A**: (i) first estimate the instant velocity 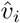 of each cell *i* = 1,…, *C*, and (ii) then estimate the non-zero entries of **A** using a stability selection procedure akin to the one used in TIGRESS to identify non-zero coefficients in the regression problem *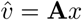* from samples 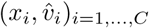. The technical details of GRISLI are presented in the Methods section.

While GRISLI involves a step of velocity inference absent from SCODE, the benefits of the GRISLI model over the SCODE model include the facts that (i) we do not need to assume that all cells lie on the same trajectory, and (ii) we make no restricting assumption on **A**, such as being of low rank and symmetric, and still derive a computationally efficient convex problem to estimate **A**.

### 3.2 Performance on GRN inference

To assess the predictive capacity and the speed of GRISLI, we test it on two benchmark datasets analyzed by Matsumoto *et al*. (2017): (i) a murine dataset of 373 cells corresponding to direct reprogramming of murine embryonic fibroblast cells to myocytes at days 0, 2, 5 and 22 (Treutlein *et al*., 2016), and a human dataset of 758 cells corresponding to differentiation of human ES cells to definitive endoderm cells, taken at 0, 12, 24, 36, 72 and 96h (Chu *et al*., 2016). We follow the experimental protocol of Matsumoto *et al*. (2017) to assess the effectiveness of GRISLI to predict known regulations, and compare it to SCODE and TIGRESS. All methods having a stochastic component, we run them 30 times on each dataset and summarize their performance by the distribution of AUC scores (see Methods), the AUC taking values between 0.5 for a random prediction to 1 for a perfect recovery of known regulations.

Figure 1 summarizes the performance of the three methods on both datasets. Since each method depends on several parameters, we tested different parameter set, as detailed in the next section, and report here the best performance of each method to assess how well they can perform if we choose good parameters. The parameters used for SCODE were the one provided for their respective datasets in Matsumoto *et al*. (2017). The parameters used for GRISLI result from an extensive search, illustrated on Figure 2. The parameters used for TIGRESS were obtained similarly, in practice they coincide with the parameters of GRISLI.

**Figure 1:**
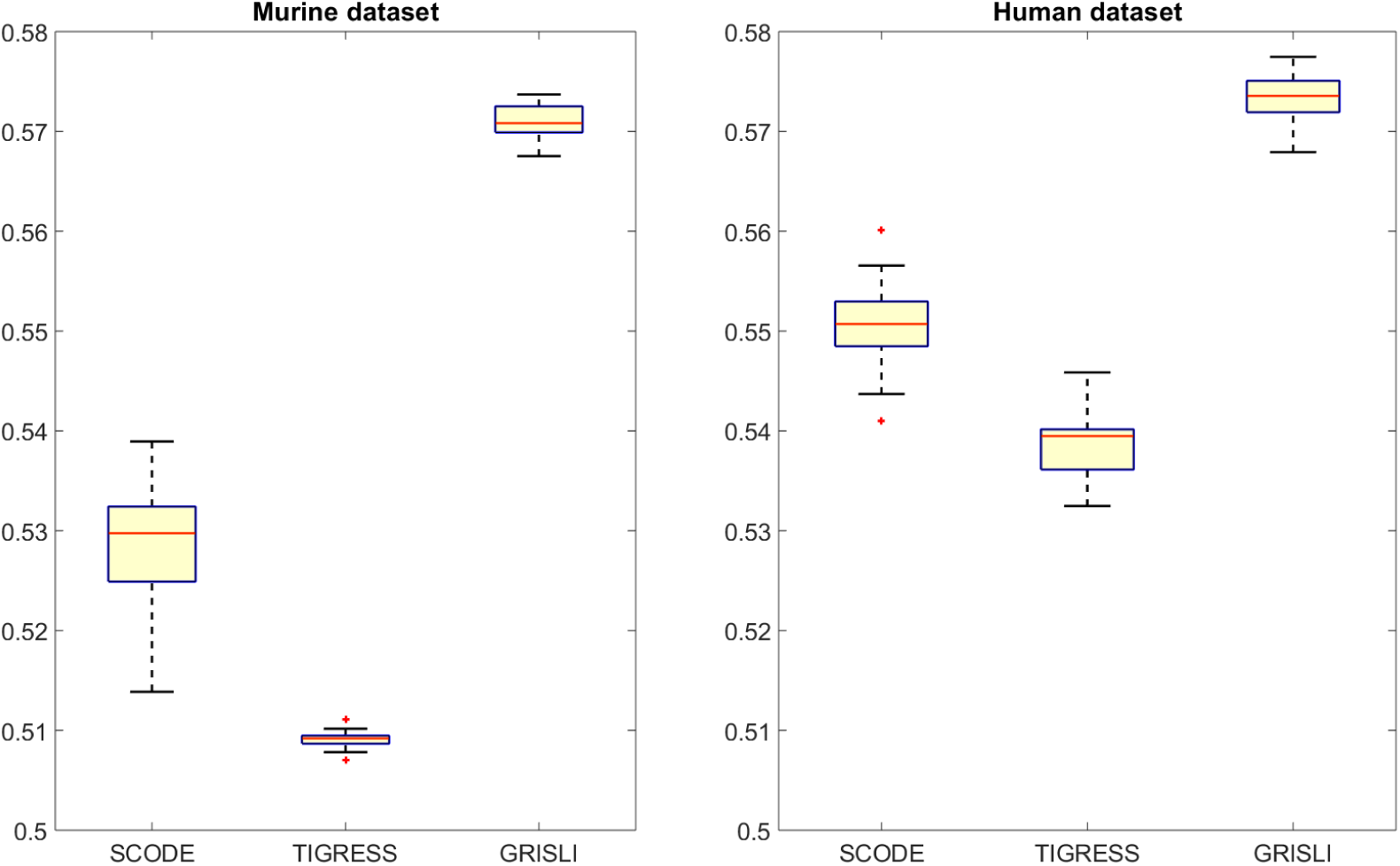
Performance of the different methods (as distribution of AUC over 30 repeats) on the murine (left) and human (right) benchmarks. SCODE score was obtained taking the average of 50 replicates with the rank *D* equal to 4 and 100 trials. GRISLI has respectively as parameters *L* = 70, *R* = 1500 and *α* = 0.3 for the murine benchmark, *L* = 1, *R* = 3000 and *α* = 0.3 for the human benchmark.TIGRESS was run with the same parameters as GRISLI.

**Figure 2:**
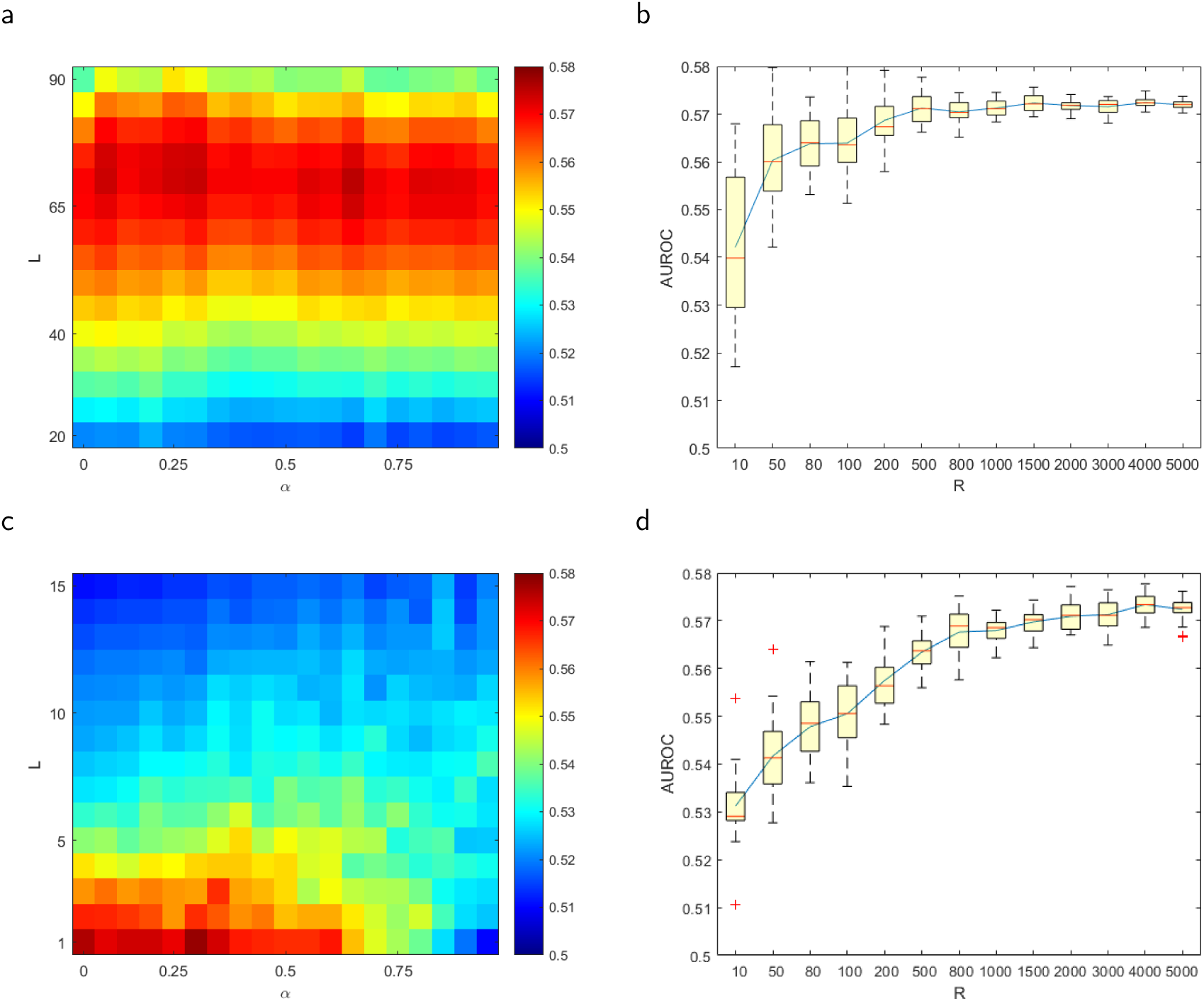
(a): Performance of GRISLI (AUC) on the murine dataset with with *R* = 1500 and varying *α* and *L*. (b): Performance of GRISLI (AUC) on the murine dataset with *α* = 0.4, *L* = 70 and varying *R* (repeated 20 times). (c): Same as (a) but for the human data and *R* = 3000. (d): Same as (b) but for the human data with *α* = 0.3 and *L* = 1.

We first notice that, in both cases, TIGRESS has the poorest performance, which highlights the limitations of GRN techniques developed for bulk RNA-seq in the context of scRNA-seq data. This is coherent with the findings of Matsumoto *et al*. (2017) who noticed for example that GENIE3, another state-of-the-art method for GRN inference from bulk RNA-seq data, performs poorly on these scRNA-seq data. Since both TIGRESS and GENIE3 are based on a steady-state assumption of expression data, this suggests that the explicit dynamical model of SCODE or GRISLI is beneficial for scRNA-seq data.

Second, we see that GRISLI outperforms SCODE on both datasets: on the murine benchmark, GRISLI has a mean AUC of 0.571 vs 0.528 for SCODE, while on the human benchmark GRISLI reaches an AUC of 0.573 vs 0.550 for SCODE. Contrary to Matsumoto *et al*. (2017), we did not provide only one AUC value to assess the performance but a boxplot that shows the inherent stochasticity of the methods. As both SCODE and GRISLI share the same underlying dynamical model, this highlights the benefits of the GRISLI approach to estimate the parameters of the system. We furthermore notice that the variability in the performance across runs is smaller for TIGRESS than for SCODE, while TIGRESS ran faster (respectively 20s and 100s on the human and murine datasets, on a 4-cores 3.5GHz Intel Core i5 with 16GB of 667MHz DDR3 RAM) than SCODE (500s on both datasets).

For the murine dataset, we used Monocle pseudo-time as a time-label. However, for the human dataset, we used the real experimental time values, as the pseudo-time AUC results did not increase with *R*. It is possible that when the measurements are close enough and evenly spaced in time (such as for the human data), real time is sufficient, while for more distant experiences (which is the case of the murine data) the pseudo-time may be of some interest.

### 3.3 Sensitivity to the parameters

Here we investigate in more details the influence of the parameters *L, α* and *R* on the performance of TIGRESS. While *R* should typically be chosen as large as possible to ensure that the empirical average converges to the the expectation in the procedure, the optimal choice of *L* and *α* is harder to predict.

We therefore systematically assess the performance of GRISLI, in terms of AUC, over a large grid of values for *L, α* and *R*. Figure 2 summarizes the results, for both the murine and the human datasets. As expected, increasing *R* is always beneficial. For example, Figures 2b and 2d show for a particular choice of *L* and *α* that, after swiftly increasing, the AUC values reaches a plateau when *R* increases. As *R* has a significant effect on runtime, we take an intermediate value of *R* = 1500 for the murine dataset and *R* = 3000 for the human dataset for the subsequent experiments.

The influence of *L* and *α* is, as expected, more complex and depends on the dataset. On the murine dataset, the AUC is fairly stable and optimal for any *L* ∈ [62, 78] with little influence of *α* (Figure 2a), while on the human dataset only the value *L* = 1 seems adequate, having a bigger effect as the scores sharply diminish for larger than 0.6 (Figure 2c). Nonetheless the overall shape is coherent with the analysis of Haury *et al*. (2012): there is a compensating effect, as a smaller *α* increases the diversity between the batches. On the contrary, reducing *L* limits the number of selected edges of the regulation network, making the predictions more similar.

The optimal values for *L* are strikingly different between the two datasets. While taking *α* equal to 0.3 or *R* larger than 1000 seem adequate in both cases, determining heuristics to choose *L* is still an open problem. In practice, we suggest to test different values of *L* and select the one that best recapitulates known interactions, if any.

## 4 Discussion and conclusion

Based on the (pseudo-)time information of scRNA-seq data, we propose GRISLI, a new method to infer GRN without any other information than the scRNA-seq data themselves. GRISLI is based on the same linear ODE formalism as SCODE, where the GRN is defined as the support of the matrix that relates TF expression to velocities of other genes; however GRISLI differs from SCODE in several assumptions. While SCODE assumes that all cells are on the same trajectory, which allows to model each cell by integrating the ODE from a unique initial condition, GRISLI considers bundles of trajectories where each cell may be following a unique trajectory, governed by a unique ODE common to all cells. Given the inherent stochasticity in gene expression, and the well-known bifurcations possible when similar cells differentiate in different subtypes (Paul *et al*., 2015), we believe that allowing cells to evolve on different trajectories is an important property. This flexibility prevents us from integrating the ODE as in SCODE, and forces us instead to estimate the local velocity of each cell. We propose a simple estimator based on a weighted average of finite differences between pairs of cells, and believe that much work remains to be done for velocity inference from scRNA-seq data. Interestingly, La Manno *et al*. (2018) proposed recently a completely different approach for velocity inference, by comparing the quantity of spliced vs unspliced mRNA; it would be interesting to assess how this estimator correlates with ours, and potentially to use it in GRISLI for GRN inference. Another difference between GRISLI and SCODE concerns the assumptions on the structure of the GRN. For computational reasons, SCODE constrains the GRN matrix to be low-rank, while GRISLI puts no assumption on it. Furthermore, if we wish to constrain the structure of the GRN in GRISLI, it can easily be achieved by adding structured sparsity constraints in the sparse regression problem (Bach *et al*., 2012).

In terms of performance, we observed that GRISLI outperforms both SCODE and TIGRESS on both human and murine scRNA-seq data. The limited performance of TIGRESS highlights the fact that methods developed for bulk RNA-seq data, based on the assumption that samples are near a steady-state condition, are not optimal for single-cell data. This was already observed by Matsumoto *et al*. (2017) with other state-of-the-art GRN inference methods for bulk RNA-seq, and confirms the relevance of interpreting the GRN as the support of the matrix that relates expression to velocity in the linear ODE framework. The fact that GRISLI outperforms SCODE, on the other hand, confirms the relevance of our assumptions and estimation procedure. However, we should keep in mind that the performance in absolute value remains modest, with a maximum AUC of 0.58. This is roughly similar to the best performances reached on simpler organisms from bulk transcriptomic data (Marbach *et al*., 2012), and highlights again the difficulty of *de novo* GRN inference. In addition, we note that GRISLI has two main parameters to tune (*L* and *α*), which have a significant impact on the performance. If some interactions are known, we suggest to tune them over a grid of candidate values by maximizing the fit between known and predicted interactions. Finding heuristics to automatically chose *L* and *α* when no known interaction is available is an interesting future work.

## Acknowledgments

The authors thank deeply Héctor Climente-González and Samyadeep Basu for enlightning discussions.

## Funding

None declared

## Conflict of Interest

None declared.

## Availability of data and materials

The code and data are available at https://github.com/PCAubin/GRISLI

